# Antibacterial activity of ethanolic and aqueous extracts of Zingiber officinale on Streptococcus pneumoniae and Pseudomonas aeruginosa

**DOI:** 10.1101/2023.01.03.522596

**Authors:** Anjellina Rukundo, Denis Omara, Samuel Majalija, Solomon Odur, Steven Alafi, Samuel George Okech

## Abstract

**Background:** *Streptococcus pneumoniae*, a capsulated lancet gram-positive bacterium, is the leading cause of mortality and morbidity among children globally and is the primary cause of pneumonia. *Pseudomonas aeruginosa* is an opportunistic human pathogen and the leading cause of nosocomial infections, among patients who are admitted to intensive care units. With the increasing resistance of microorganisms to antibiotics, there is a shift of choice from allopathy to naturopathy, where herbs are common ingredients of medicines and components of treatment protocols. It is against this background that this study aimed to investigate the susceptibility of *P. aeruginosa* and *S. pneumoniae* to ethanolic and aqueous extracts of ginger using the agar well diffusion technique.

**Methodology:** Absolute (95%) Ethanol and distilled water were used as solvents to make extracts from the ginger powder. The filtrate was dried, and the resulting substance was used to conduct antimicrobial tests on *Streptococcus pneumoniae* and *Pseudomonas aeruginosa* isolates using the agar well diffusion technique. The diameters of inhibition zones were measured, and statistical analysis was done by one-way ANOVA. Minimum inhibitory and bactericidal concentrations were determined by serial dilution.

Freshly prepared sterile distilled water was used as negative control and ciprofloxacin (5 *μ*g/disk), an antibiotic was used as positive control.

**Results:** The test organisms were sensitive to both ethanolic and aqueous extracts of ginger. However, this was highly dependent on the concentrations of the extracts. The ethanolic extract had lower Minimum Inhibitory and Bactericidal Concentrations than aqueous extract on both bacterial species and at a concentration of 2g/ml, the ethanolic extract was 2-fold and 1.6-fold more effective in inhibiting the growth of *P. aeruginosa* and *S. pneumoniae* respectively than aqueous extract.

**Conclusion:** With the high susceptibility of the tested bacterial isolates to the ginger extracts used in this study, it is evident that ginger extracts can serve as a suitable antibiotic. However, further studies need to be conducted on the antimicrobial effects of ginger extracts on other bacterial species.

## 1. Introduction

*Streptococcus pneumoniae*, a capsulated lancet gram-positive bacterium, is the leading cause of mortality and morbidity among children globally and is the major cause of pneu monia [1]. World Health Organization estimated that 467,000 deaths among children under five years are caused by Pneumococcal infections. Disease rates and mortality are higher in developing countries with majority of deaths occurring in Africa[2]. Antibiotic treatment of Pneumococcal disease is being compromised by the world wide emergence of drug resistant strains[3]. *Pseudomonas aeruginosa* also being an opportunistic human pathogen, is the leading cause of nosocomial infections, especially among patients who are admitted to intensive care units (ICU) and it has been implicated in diverse nosocomial infections like nosocomial pneumonias, urinary tract infections (UTIs), skin and soft tissue infections, in severe burns and in infections in immuno-compromised individuals [1] [4] [5]. In recent years, a considerable increase in the prevalence of multidrug resistance (MDR) in *P. aeruginosa* has been noticed, which is related to high morbidity and mortality [6].

Perhaps, this high degree of multidrug resistance is related to the presence of antibiotic efflux systems which provide resistance to multiple antimicrobial agents [7]. *Pseudomonas aeruginosa* is frequently resistant to many commonly used antibiotics [8].

With increasing resistance of microorganisms to antibiotics, there is a shift of choice from allopathic to ayurvedic and naturopathic practices, where herbs and spices are very common ingredients of medicines [9]. Medicinal plants can be used to combat antimicrobial resistance. The antimicrobial resistance (AMR) crisis is the increasing global incidence of infectious diseases affecting the human population, which are untreatable with any known antimicrobial agent [10]. With increasing use of drugs, microorganisms are attaining resistance to commonly used antibiotics, which leads to downfall of effectiveness of conventional medicines and therefore, search for new antimicrobial agents has become necessary [11].

Medicinal plants have a long history of use for the benefit of mankind [12]. About 80% of the world’s populations rely mainly on traditional therapies which involve the use of plant extracts or their active substances according to the report of the World Health Organization [13]. Medicinal plants play an important role in traditional health care systems for curing many diseases. The medicinal value of these plants lies in some chemical substances that produce a definite physiological action on human body [14]. Plants have been used as a source of therapeutic agents in traditional medicinal system since ancient times due to bioactive compounds they contain [15]. World Health Organization has described traditional medicine as a cheap way to achieve total health care coverage of the world’s population and has encouraged rational use of the plant-based traditional medicine by her member states[16]. Herbs are generally considered safe and proved to be effective against various human ailments and their medicinal uses have been gradually increasing in developed countries [17]. Products derived from plants have been used for medicinal purposes for centuries and at present about 80% of the world population relies on botanical preparation as medicines to meet their health needs [18]. Traditional medicines have been used for many centuries by a substantial proportion of the population of India [19]. The interest in the study of medicinal plants as a source of pharmacologically active compounds has increased worldwide [20]. The failure of antibiotics has resulted in the search for more effective sources of natural products from plants and they have been found safe and good source of pharmacological effect for man [7].

In recent studies, a few gingerol-related components have been found to possess antibacterial and antifungal properties [21]. Ginger derives its name from the genus Zingiber and the family Zingiberaceae [22]. Zingiberaceae is commonly grown for its edible rhizome which is widely used as a spice and medicine [23]. Besides its use as a spice, ginger is also a medicinal plant that has been widely used all over the world, for treatment of a wide array of unrelated ailments including arthritis, cramps, rheumatism, sprains, sore throats, fever and infectious disease [20]. It has direct antimicrobial activity and thus can be used for treatment of bacterial infections [24]. The volatile oil, gingerol is the most medically powerful because it inhibits prostaglandin and leukotriene formation, which are products that influence blood flow and inflammation[2]. It has direct antimicrobial activity thus can be used for treatment of bacterial infections and its antimicrobial and other biological activities are attributed to gingerols, shogaols and zingerone [25]. In particular, its gingerol-related components have been reported to possess antimicrobial and antifungal properties, as well as several pharmaceutical properties [21].

The herb is generally considered safe and proved to be effective against human ailments and their medicinal uses have been gradually increasing in developed countries [26]. It has been shown to be effective on ischemia/reperfusion (I/R) injury in the rat’s kidney [27]. It has long been used as a naturopathy due to its potential antimicrobial activity against different microbial pathogens [24]. It has several ethno-medicinal and nutritional values as a spice and flavoring agents in Ethiopia and elsewhere. In the last few decades, ginger was extensively studied for its medicinal properties by advanced scientific techniques [28]. It is the most effective remedy against nausea and vomiting associated with surgery [29]. In laboratory animals, gingerols increase the motility of gastro intestinal tract and have analgesic, sedative, antipyretic and antibacterial properties [30]. However, the antibacterial activity of ginger against the drug resistant organisms like *Streptococcus pneumoniae* and *Pseudomonas aeruginosa* has been scarcely explored therefore there is a perpetual need to exploit its effect on these multi-drug resistant organisms [31].

## 2. Methodology

### 2.1. Research design

This was an experimental study which was carried out in Pharmacology Laboratory and Microbiology Laboratory, College of Veterinary Medicine, Animal Resources and Biosecurity, Makerere University - Uganda. The ginger was bought randomly from the sellers in Kasubi market during the month of January 2019 and was put in sterile bags and transported to the Microbiology Laboratory.

### 2.2 Sample collection

The ginger rhizome was bought from Kasubi market which is one of the biggest local markets in Kampala, Uganda. The rhizomes were packaged in sterile bags and delivered to the microbiology laboratory.

### 2.3 Laboratory methods

#### 2.3.1 Preparation of ginger extract

Ginger rhizomes were washed with clean water and rinsed several times in sterile distilled water. They were sliced into pieces using a sterile knife, air dried for three weeks and ground into powder using a grinder. One hundred sixty (160g) grams of smooth powder were obtained. The Ethanolic and Aqueous extracts were prepared by dissolving 80g of ginger powder in 400mls of each solvent (i.e., 95% Ethanol and distilled water) in amber bottles and the mixtures were left for 4 days while agitating them daily for uniform extraction to occur. The suspension was filtered using Whatman filter paper. The filtrates were evaporated to dryness in a water bath at a temperature of 55 *°*C and preserved for the antimicrobial activity test. The dry extracts were dissolved in dimethyl sulphoxide (DMSO) – (SIGMA D2650, Germany) to allow complete solubilization. Two different concentrations of each extract were used i.e. (0.5 g/ml and 2 g/ml.) and 10% DMSO was used as the solvent.

#### 2.3.2 Preparation of bacterial cultures

Standard cultures of Streptococcus pneumoniae and *Pseudomonas aeruginosa* were obtained from the Microbiology laboratory at the College of Veterinary Medicine, Animal Resources and Biosecurity, Makerere University - Uganda. The pure laboratory isolates were resuscitated by emulsifying one loopful of organisms in 1ml of normal saline. The resuscitated isolates of *S. pneumoniae* and *P. aeruginosa* were inoculated using streaking method of inoculation on the surface of King’s agar (Cat no: M1544F-500G, Himedia Biosciences, India) and MacConkey agar (Cat no: MH081-500G, Himedia Biosciences, India) respectively. These culture media were prepared according to the manufacturer’s instructions. The inoculated plates were incubated for 24 hours at 37^0^c to obtain fresh colonies.

### 2.4 Assessment of antibacterial activity

#### 2.4.1 Agar well diffusion

Fresh Colonies of the microorganisms obtained, were emulsified in normal saline and the turbidity adjusted by comparing with that of 0.5 McFarland standard of Barium sulphate solution (equivalent to 1×10^6^ CFU/mL) until the turbidities were the same. Bacterial lawns of the organisms were prepared on the sterile Muller Hinton Agar (Cat no: M173-500G, Himedia Biosciences, India) plates by inoculating the microorganisms evenly on the surface of the media by streaking plating method. The prepared bacterial lawns were then left for 15 minutes at room temperature and six (6) holes were made on the agar plate using sterilized cork-borer. Both ethanolic (0.1 ml) and aqueous (0.1 ml) extracts of each concentration (0.5g/ml and 2g/ml) were dispensed into respective wells and 0.1 ml of freshly prepared sterile distilled water was used as a negative control. Ciprofloxacin (5 *μ*g/disk), an antibiotic was used as a positive control.

All the two extracts were applied against *P. aeruginosa* and *S. pneumoniae* at two different concentrations (0.5g/ml and 2g/ml). Thirty (30) minutes pre-diffusion time was allowed, after which the plates were incubated at 37*°*C for 24 hours. The antimicrobial agent diffused in the agar medium and inhibited the growth of the bacteria being tested. The zones of inhibition were then measured using the zone of inhibition ruler. Each test was carried out in 8 replicates to ensure quality and the mean of the diameters of the resulting inhibition zones were recorded in millimeters.

### 2.5 Determining minimum inhibitory concentrations (MIC and Minimum Bactericidal Concentrations (MBC)

The minimum inhibitory concentration (MIC) of the ginger extract against the test organisms was determined using 2-fold serial dilution. Fresh culture plates of the microorganisms were prepared by dissolving them in normal saline. Five hundred microliters (500*μ*l) of Mueller Hinton Broth (Cat no: M173-500G, Himedia Biosciences, India) were put in a set of the six tubes each, which were then plugged with cotton wool at the end before being sterilized by autoclaving. After sterilization, 500*μ*l of the extract (Ethanolic and Aqueous) was put in the first tube to make 1 ml and thereafter 500*μ*l was picked from tube one and transferred to tube two and the same was done for tube three to tube six. On the last tube six, 500*μ*l was discarded off. The resulting concentrations for tube 1, 2, 3,4, 5 and 6 were 5g/ml, 2.5g/ml, 1.25g/ml, 0.625g/ml, 0.313g/ml and 0.156g/ml respectively. In each of the six tubes, 0.1ml of the microorganisms were added and the tubes were incubated overnight. The tubes with the lowest concentration of the extract showing no turbidity (no bacterial growth) were reported in this study as the MIC. After incubation, Minimum Bactericidal Concentrations (MBC) were determined by taking a loopful from each test tube and sub-culturing it on Mueller Hinton Agar to see if bacterial growth was inhibited. From this, the minimum concentration of extract that was bactericidal was determined.

### 2.6 Statistical analysis of ZOI results

The data obtained was entered in Microsoft Excel and analyzed using IBM SPSS. One-way ANOVA was used to determine whether there was a significant variation in the ZOI for the different fractions of extracts that the test organisms were subjected to.

## 3. Results

### 3.1 Antibacterial activity of ginger extracts

The test organisms were sensitive to both ethanolic and aqueous extracts of ginger. However, this was highly dependent on the concentrations of the extracts. The aqueous extract was not effective on *P. aeruginosa* at lower doses (0.5g/ml) while at a higher dose (2g/ml), it showed activity against *P. aeruginosa* (Table 1). The ethanolic extract was very effective against all the test microorganisms both at lower and higher concentrations with the Zone of Inhibition increasing with increase in concentration (Table 1 and Figure 1).

**Figure 1:**
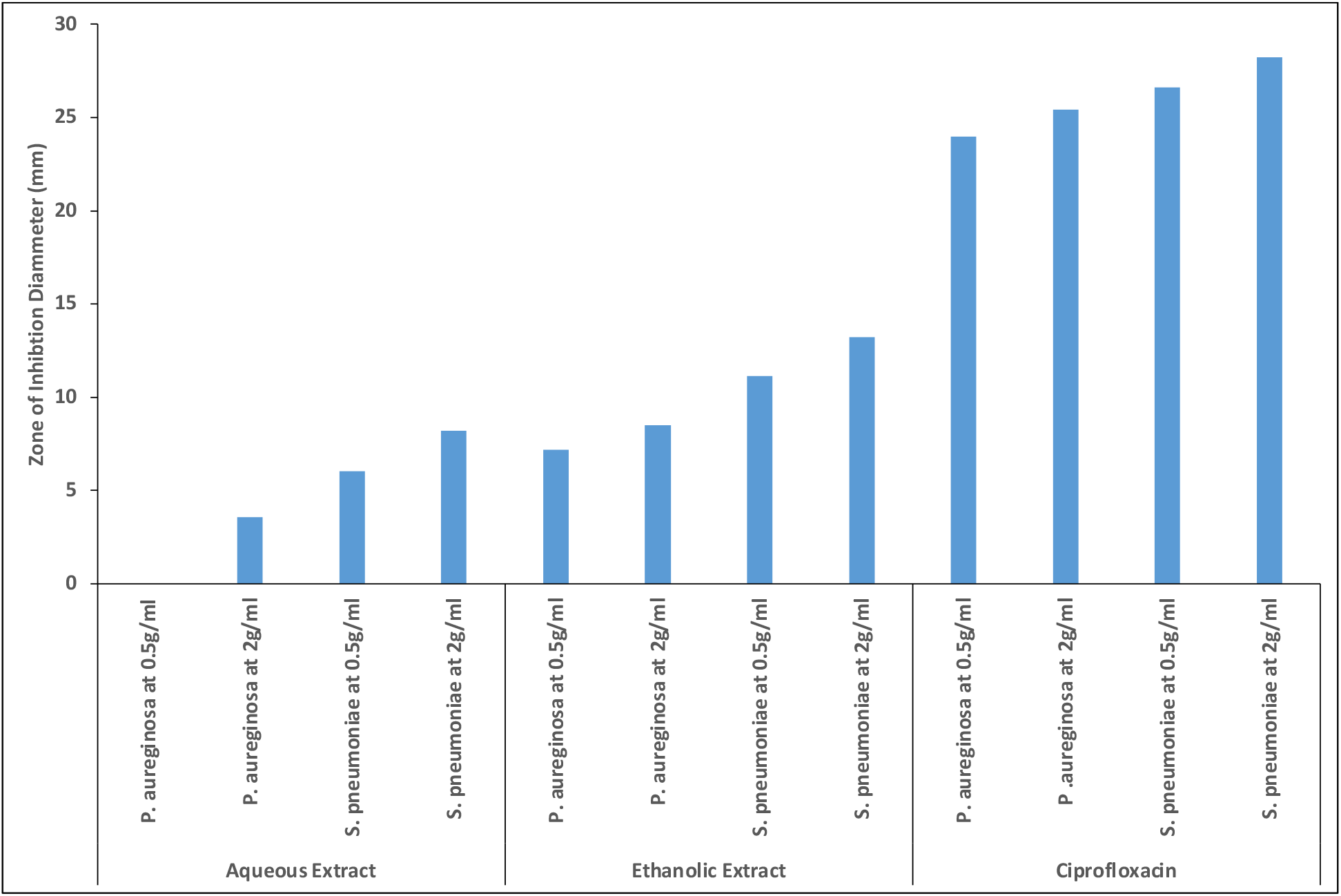
A graph showing zone of inhibition of ethanolic and aqueous extracts of ginger

**Table 1:**
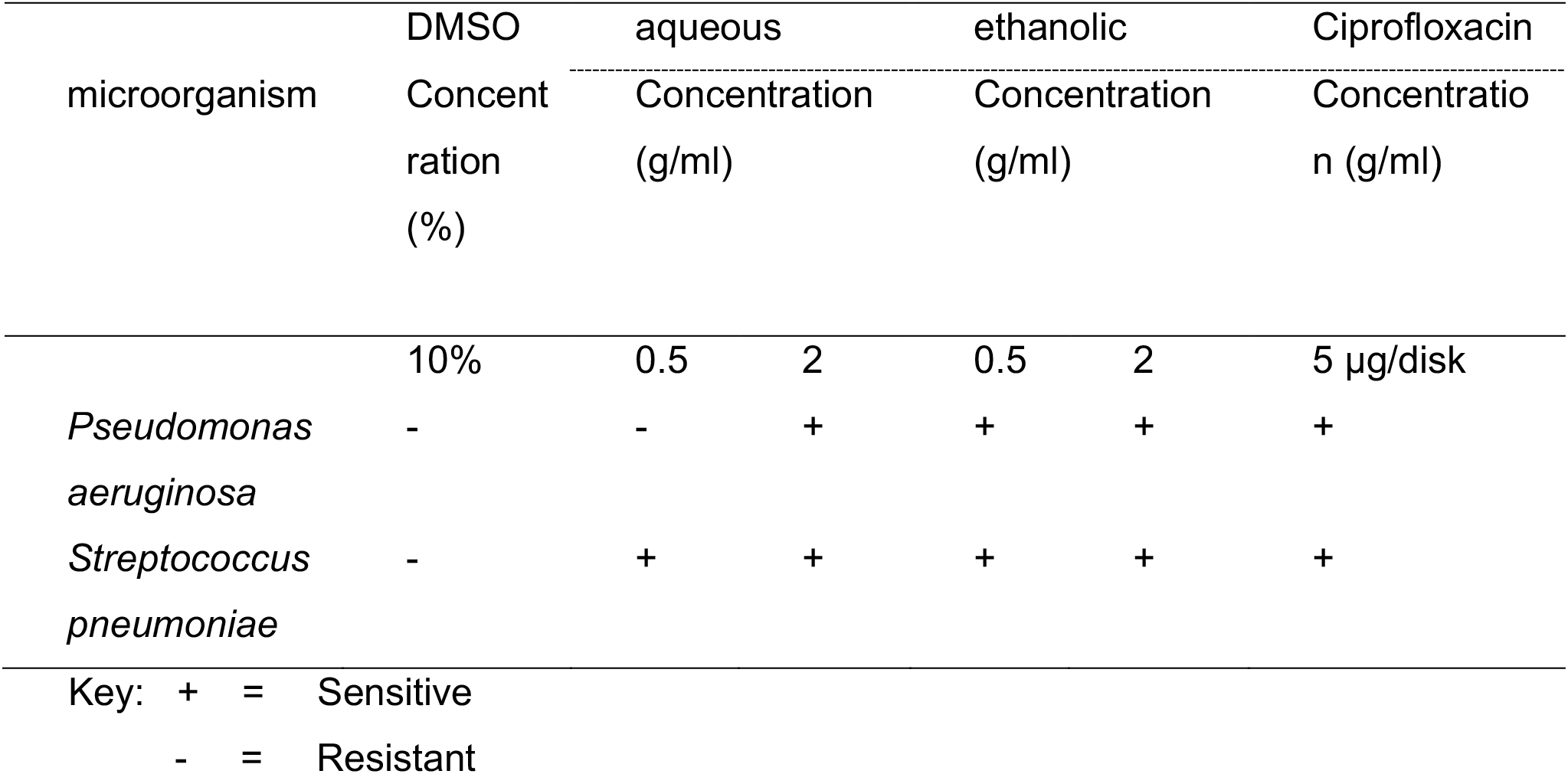
The antibacterial activity of ethanolic and aqueous extracts of Ginger on *P. aeruginosa* and *S. pneumoniae*.

At a concentration of 2g/ml, ethanolic extract of ginger was 2-folds and 1.6-folds more effective in inhibiting growth of *P. aeruginosa* and *S. pneumoniae* respectively than aqueous extract of ginger and from the results of one-way ANOVA, the P-values obtained were all less than 0.05 at 95% confidence interval for each of the test organisms (Figure 1). Organic solvent DMSO used was used as negative control and standard drug (ciprofloxacin) was far most effective than the tested extracts having highest zone of inhibition (Figure 1).

### 3.2 Minimum Inhibitory Concentration (MIC) and Minimum Bactericidal Concentrations (MBC) of ethanolic and aqueous extracts of ginger

The Minimum Inhibitory Concentration and Minimum Bactericidal Concentrations (MBC) of both aqueous and ethanolic extracts against the two test organisms were determined as shown in Table 2. The MIC and MBC of aqueous extract against *P. aeruginosa* was not determined since it had not shown activity at a concentration of 0.5g/ml. The growth of *P. aeruginosa* was inhibited at the lowest concentration of 1.25 g/ml (MIC) of ethanolic extract and killed at the lowest concentration of 2.5 g/ml of ethanolic extract (MBC) after 18 hours and these were the same for S. pneumoniae. The growth of *S. pneumoniae* was inhibited at 2.5g/ml of the aqueous extract and killed at 5g/ml of the aqueous extract whereas for the ethanolic extract the growth of *S. pneumoniae* was inhibited at 1.25g/ml and killed at 2.5g/ml. This shows that ethanolic extract was relatively more effective than the aqueous extract.

**Table 2:**
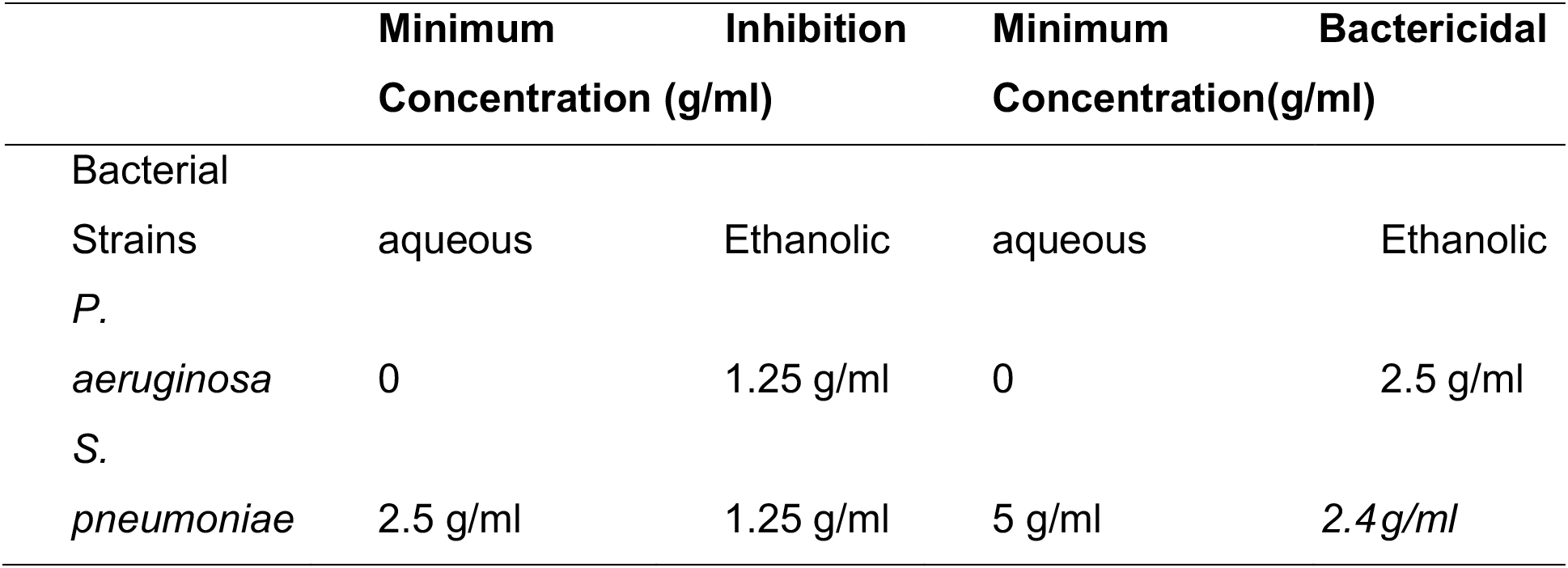
Minimum inhibitory concentrations of ethanolic and aqueous extracts of ginger.

## 4. Discussion

In this study, both ethanolic and aqueous extracts demonstrated antibacterial activity against *P. aeruginosa* and S. pneumoniae. The antibacterial activity of ginger extracts could be attributed to the chemical properties of ginger. Phytochemical investigation of several types of ginger rhizomes has indicated the presence of bioactive compounds, such as gingerols, which are antibacterial agents and shogaols, phenylbutenoids, diarylheptanoids, flavanoids, diterpenoids, and sesquiterpenoids, thus the bioactive ingredients were responsible for the antibacterial activity of the plant extracts [32]. However, inhibition of bacterial growth was highly dependent on the concentration of the extracts where the antimicrobial activity of the extracts was higher at high concentration and lower or inactive at very low concentrations. Thus, the study may suggest that the inhibition of bacterial growth by the extracts is dose dependent. The ethanolic extract showed more antibacterial activity than aqueous extract. The solvent of extraction and method could have affected the degree of antibacterial activity. The stronger extraction capacity of ethanol could have produced greater amount of the active constituents responsible for the higher antibacterial activity.

The Minimum Inhibitory Concentrations (MICs) and Minimum Bactericidal Concentrations (MBCs) of the Aqueous extract were two-folds higher than those of the Ethanolic extract. This could be attributed to the fact that concentration of the bioactive compounds in ethanolic extracts were relatively higher than they were in the aqueous extract. Gingerol, a bioactive compound in ginger extracts is a phenol phytochemical compound which is soluble in organic solvents like ethanolic, isopropanol, methanol, among others whereas in inorganic solvents like water, it is insoluble [33] [34] [35] [36]. Therefore, the solubility factor could have led to extraction of higher yield of gingerol in ethanol than in distilled water [37] hence accounting for the lower MICs of the ethanol extract than the aqueous extract.

*S. pneumoniae* was more susceptible to extracts than *P. aeruginosa*. This could be attributed to the fact that gram-positive bacteria are known to be more sensitive to plant extracts than the gram negative [38]. These observations are likely to be a result of the differences in the cell wall structure between gram negatives and gram positives, with gram negative outer membrane acting as a barrier to many environmental substances, including antibiotics [39]. The outer membrane, a characteristic of gram-negative bacteria like *P. aeruginosa* functions as a molecular sieve through which molecules with molecular mass >600–1000 Da cannot penetrate [38].

Despite the presence of porins with low specificity, the outer membrane shows very low permeability toward hydrophobic compounds, which has been ascribed to the presence of the lipophilic LPS [40] [41]. Gingerol, the bio-active compound in the extracts is hydrophobic [42] and this could also explain the resistance shown by *P. aeruginosa* to the lower concentrations of the extract as compared to S. pneumoniae. Another unique feature of *P. aeruginosa* is its resistance to a variety of antibiotics which is attributed to a low permeability of the cell wall, the production of inducible cephalosporinases, an active efflux and a poor affinity for the target (DNA gyrase) [39].

## 5. Conclusions

This study has shown that both *S. pneumoniae* and *P. aeruginosa* were susceptible to aqueous and ethanolic extracts of ginger in vitro hence affirming the presence of antibacterial compounds in ginger rhizomes. *S. pneumoniae* was more susceptible to both extracts than *P. aeruginosa*. However, inhibition of bacterial growth was highly dependent on the concentration of the extracts where the antimicrobial activity of the extracts was higher at high concentration and lower or inactive at very low concentrations. With the high susceptibility of the tested bacterial isolates to the ginger extracts used in this study, it is evident that ethanolic and aqueous extracts of ginger contain compounds that can serve as suitable antimicrobial chemotherapeutics against *S. Pneumoniae* and *P. aeruginosa*. However, further studies need to be conducted to assess the safety of the extracts especially their cytotoxicity, and identification of the specific compounds with antimicrobial activity. The antimicrobial effects of the ginger extracts on other bacterial species that have shown resistance to most of the antibiotics routinely used in treatment of bacterial infections as well as on fungi should also be studied.

## Acknowledgements

This study was funded by MasterCard Foundation through its Scholars Program at Makerere University. We acknowledge the Microbiology laboratory and entire team at College of Veterinary Medicine, Animal Resources and Biosecurity (COVAB), Makerere University, Uganda, whose efforts were invaluable.

## Authors’ Contributions

All authors had equal contributions to all stages of this study and all authors read and approved the final manuscript. The funders had no role in the design of the study; in the collection, analyses, or interpretation of data; in the writing of the manuscript, or in the decision to publish the results.

